# Asymmetry in catalysis by *Thermotoga maritima* membrane-bound pyrophosphatase demonstrated by a non-phosphorous allosteric inhibitor

**DOI:** 10.1101/423954

**Authors:** Keni Vidilaseris, Alexandros Kiriazis, Ainoleena Turku, Ayman Khattab, Niklas G. Johansson, Teppo O. Leino, Paula S. Kiuru, Gustav Boije af Gennäs, Seppo Meri, Jari Yli-Kauhaluoma, Henri Xhaard, Adrian Goldman

**Author notes:** Correspondence and requests for materials should be addressed to A.G.

## Abstract

Membrane-bound pyrophosphatases are homodimeric integral membrane proteins that hydrolyse pyrophosphate into orthophosphates, coupled to the active transport of protons or sodium ions across membranes. They are important in the life cycle of bacteria, archaea, plants, and protist parasites, but no homologous proteins exist in vertebrates, making them a promising drug target. Here, we report the first non-phosphorous allosteric inhibitor (*K*_i_ of 1.8 ± 0.3 μM) of the thermophilic bacterium *Thermotoga maritima* membrane-bound pyrophosphatase and its bound structure at 3.7 Å resolution together with the substrate analogue imidodiphosphate. The unit cell contains two protein homodimers, each binding a single inhibitor dimer near the exit channel, creating a hydrophobic clamp that inhibits the movement of β-strand 1–2 during pumping, and thus preventing the hydrophobic gate from opening. This asymmetry of inhibitor binding with respect to each homodimer provide the first clear demonstration of asymmetry in the catalytic cycle of membrane-bound pyrophosphatases.

## Introduction

Membrane-bound pyrophosphatases (mPPases) are a family of enzymes that hydrolyse pyrophosphate into two phosphates and couple this with proton and/or sodium transport across the membrane, thus creating an electrochemical gradient. These enzymes were initially discovered in photosynthetic bacteria and plants^1,2,3^. They were later found in protist parasites, archaea, and certain species of bacteria, but do not occur in animals and humans^4,5^. In various organisms, these proteins are important for the survival under diverse stress situations due to energy limitation, such as osmotic stress, anoxia, mineral deficiency, low temperature, and intense light^6^. In bacteria and archaea, mPPases reside in the cell membrane^4^, while in protists, algae and plants, the proteins can also be located in acidocalcisomes, the vacuole, and/or Golgi apparatus^7,8^.

mPPases are homodimeric. Each monomer contains 15–17 transmembrane helices with a molecular weight of 70–81 kDa. So far, there are structures of only two mPPases: four from *Thermotoga maritima* (TmPPase) and two for mung bean (*Vigna radiata*: VrPPase). TmPPase structure has been solved in the resting state (TmPPase:Ca:Mg)^9^, with two phosphates bound (TmPPase:2P_i_)^9^, with the substrate-analogue imidodiphosphate (IDP) bound (TmPPase:IDP)^10^, and with the phosphate analogue (WO_4_)-bound (TmPPase: WO_4_)^10^. VrPPase has been solved in the IDP-bound (VrPPase:IDP)^11^ and single phosphate-bound states (VrPPase:P_i_)^10^ In all the structures, the mPPase is a symmetric homodimer with each monomer consisting of 16 transmembrane helices (TMHs). These helices form two concentric layers with six helices (TMH5–6, 11–12, and 15–16) forming the inner layer and the other ten (TMH 1–4, 7–10, and 13–14) forming the outer layer. Each monomer consists of four regions: a hydrolytic centre, a coupling funnel, an ion gate, and an exit channel^12^ (Fig. 1a). The hydrolytic centre and coupling funnel are comprised of TMH5–6, 11–12, and 15–16 (Fig. 1b). Upon binding of substrate, the active site is closed by a long loop between TMH5 and TMH6 and opened again after hydrolysis and ion pumping^9,10,11^. In the structure of TmPPase:IDP, the Na^+^ binds within the membrane plane at the ionic gate between TMH6 and TMH16. Na^+^ is pentacoordinated by the carboxylate groups of D243^6.50^, E246^6.53^, and D703^16.46^, the O^⍰^ of S247^6.54^ and with the main-chain carbonyl of D243^6.50 10^ (Fig. 1c). (Residues are numbered as in ^12^, shown as XΣ*^a.b^* where X is the amino acid, Σ is amino acid position in TmPPase, *a* is the helix number and *b* is the amino acid position according to a central conserved residue in each helix).

**Figure 1.**
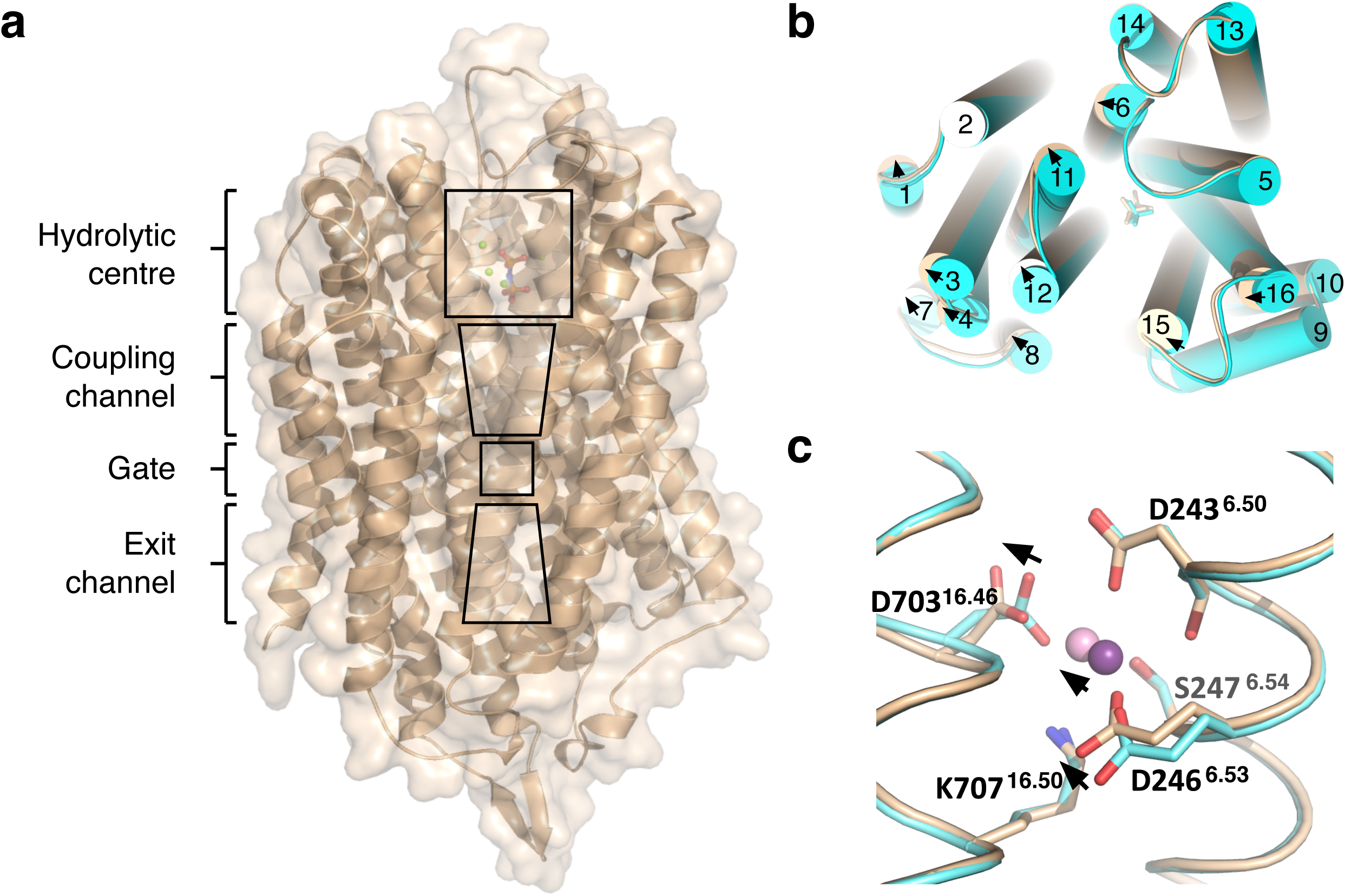
Overview of TmPPase structure. (**a**) Monomer showing the location of the hydrolytic centre, coupling channel, ion gate, and the exit channel (**b**) Top view of the superposition of TmPPase:IDP:ATC (wheat) and TmPPase:IDP complex (cyan) structure showing relative TMHs movement (arrow) upon the binding of ATC. (**c**) Superposition of the gate region between two structures (TmPPase:IDP:ATC (wheat) and TmPPase:IDP complex (cyan)). D246^6.53^, D703^16.46^, and Na^+^ slightly moving away (arrow) at the same direction relative to their positions in TmPPase:IDP structure. Violet-purple coloured sphere is the Na^+^ of TmPPase:IDP structrure.

Protist parasites such as *Plasmodium falciparum, Toxoplasma gondii, Trypanosoma brucei*, and *Leishmania donovani* all possess mPPases^13^, two types (H^+^-pump K^+^-dependent and H^+^-pump K^+^-independent mPPase) for Plasmodium^14^ and one type (H^+^-pump K^+^-dependent) for the others^15,16,17^. The mPPases, along with a V-ATPase, maintain the ionic gradient across the acidocalcisome membrane, which is necessary for acidocalcisome function, such as osmotic homeostasis upon passage of the parasite from the insect vector into the mammalian bloodstream^8^. Indeed, knockdown/knockout of mPPases causes severe reduction in polyphosphate and the loss of acidocalcisome acidity, leading to the failure of the parasites to stabilize their intracellular pH upon exposure to external basic pH^18^. In addition, it has been shown that mPPase is essential for virulence of *T. gondii* in a mouse model^19^, and both mPPases are essential genes in *P. falciparum* (Totanes, *unpubl.*). mPPases are thus promising targets for structure-based drug design because they are essential, do not exist in multicellular animals, and structures in different conformations are known.

Non-hydrolysable PP_i_ analogues are not viable mPPase drug candidates, as they will inhibit the many human enzymes that produce or hydrolyse PP_i_, from inorganic pyrophosphatases to polymerases and ectonucleotide pyrophosphatase/phosphodiesterases^20^. Our goal is thus the discovery of drug-like small molecular weight compounds that would specifically inhibit mPPases from protist parasites^13^. Here, we report the first non-phosphorous non-substrate analogue low micromolar inhibitor of TmPPase, *N*-[(2-amino-6-benzothiazolyl)methyl]-1*H*-indole-2-carboxamide (ATC), its preliminary structure-activity relationships, and a 3.7 Å resolution structure of ATC bound to TmPPase. The binding site of ATC is allosteric and furthermore asymmetric with respect to the structural homodimer, providing strong evidence that the substrate binding and catalysis events process asymmetrically.

## Results

### Structure overview

We solved the 3.7 Å resolution of TmPPase structure in complex with IDP and ATC (TmPPase:IDP:ATC) by molecular replacement using the TmPPase:IDP structure (PDB ID: 5LZQ)^10^ as the search model but with the IDP removed. Molecular replacement identified four TmPPase molecules (two dimers) in the asymmetric unit. After the first round of refinement, we observed positive F_o_–F_c_ density at 3.0 σ at the hydrolytic centre (Supplementary Fig. 1a) and at the ionic gate (Supplementary Fig. 1b) in all four monomers. We built the Mg_5_IDP complex into the former and a sodium ion into the latter. We also observed positive 3.0 σ F_o_–F_c_ density near the exit channels of chains A and C that could be fit with ATC molecules. Two molecules of ATC (ATC**-1** and ATC**-2**) could be fit at the interface between A and D (Fig. 2a and Supplementary Fig. 1c), but only one (ATC**-3**) between B and C, as the density there was weaker (data not shown). After refinement, manual rebuilding and manual placement of the IDP and ATC into the electron density, we were able to refine it to an R_work_/R_free_ of 22.5%/26.8% with acceptable stereochemistry (Table 1). Unlike previously solved TmPPase structures^9,10^, there are four molecules (two dimers, AB and CD) in the asymmetric unit (Supplementary Fig. 2a). The root mean squared deviation (RMSD) per Cα is 0.30 Å^2^ for the AB versus CD dimer, and 0.245–0.304 Å^2^ when comparing individual monomers (Supplementary Table 1).

**Figure 2.**
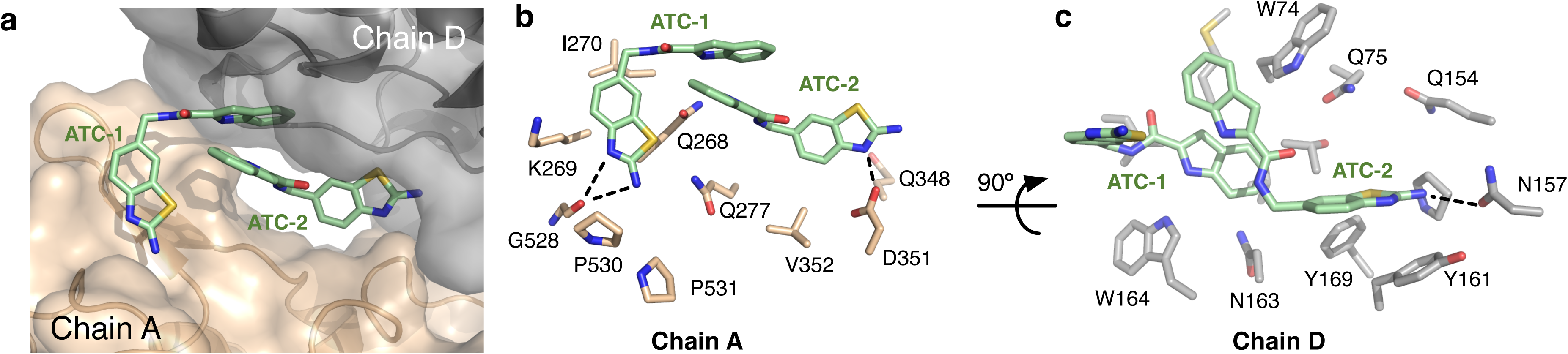
ATC interactions with TmPPase domains. (**a**) ATC located on the interface of chain A (wheat) and D (grey) in a hydrophobic cleft. Pale-green: carbon, blue: nitrogen, red: oxygen and yellow: sulphur. (**b**) Residues in chain A that are important for the interactions with the ATC dimer. (**c**) Residues on chain D that are important for the interactions with the ATC dimer. Dashed lines represent hydrogen bonds.

**Table 1.** X-ray data collection and refinement statistics.

| Data Collection |  |
| --- | --- |
| Space group | P 2<sub>1</sub>2<sub>1</sub>2<sub>1</sub> |
| Cell dimensions |  |
| *a*, *b*, *c* (Å) | 98.133, 141.63, 252.06 |
| Source | ESRF - ID29 |
| Wavelength (Å) | 0.9677 |
| Resolution (Å) | 47.41 – 3.7 (3.832 – 3.7) |
| R-merge | 0.1201 (1.451) |
| *I*/σ(*I*) | 9.90 (1.24) |
| Completeness (%) | 99.35 (99.95) |
| Multiplicity | 5.3 (5.6) |

| Refinement |  |
| --- | --- |
| Resolution (Å) | 47.41 – 3.7 |
| No. of total reflections | 203154 (21201) |
| No. of unique reflections | 38085 (3794) |
| R<sub>work</sub>(%)/R<sub>free</sub>(%) | 22.5/26.8 |
| No. of atoms | 21075 |
| Protein | 20929 |
| ligands | 145 |
| No. of chains (ASU) | 4 |
| B-factors (Å<sup>2</sup>) |  |
| Protein | 134.56 |
| Ligands/Ion | 144.93 |
| R.m.s. deviations |  |
| Bond lengths (Å) | 0.004 |
| Bond angle (°) | 0.62 |
| Ramachandran statistics |  |
| Favored (%) | 94.12 |
| Allowed (%) | 5.71 |
| Outliers (%) | 0.17 |

Statistics for the highest-resolution shell are shown in parentheses.

There are thus two dimer interfaces in the crystal structure: the AB dimer described before^9,10^ and ATC-mediated AD and BC dimers (Supplementary Fig. 2). Are these latter dimers physiologically relevant? It is fairly clear that the answer is no. First, the interactions are weak for a protein interface (219 Å^2^ buried) and the proteins interact tail to tail (Supplementary Fig. 2b), placing the extracellular loops together in a manner that would not allow a bilayer to form. Second, size-exclusion chromatography-multiangle laser light scattering (SEC-MALLS) analysis showed that the addition of ATC to the TmPPase:IDP complex (158.9 ± 0.1 kDa) did not change the oligomerisation state of the protein (162.8 ± 3.7 kDa for TmPPase:IDP:ATC complex) (Supplementary Fig. 3). ATC clearly interacts as a dimer, binding to both the E_2_S and E_2_S_2_ complexes in solution (Table 4). We have shown this both structurally and functionally, as we describe below.

The overall structure of TmPPase:IDP:ATC complex is very similar to the structure of the TmPPase:IDP complex with an RMSD/Cα of 0.46 Å and 0.42 Å to dimers AB and CD, respectively. (In what follows, all structural alignments were made with respect to monomer A unless otherwise stated.) Even though the volume of the hydrolytic pocket in TmPPase:IDP:ATC is very similar to that of TmPPase:IDP (707 Å^3^ *versus* 720 Å^3^), based on the structural alignment on TMH13–14 (residues 542–629), there are some slight clockwise movements on the cytoplasmic side of the protein helices relative to the TmPPase:IDP structure (Fig. 1b). This causes a slight movement of the Mg_5_IDP complex in the same direction as the helices. In addition, at the gate of the TmPPase:IDP:ATC structure, Na^+^ is tetracoordinated by D243^6.50^, D246^6.53^, D703^16.46^ and S247^6.54^ (Fig. 1c). This is in contrast to TmPPase:IDP, where Na^+^ appears to be pentacoordinated, with the fifth ligand being the main-chain carbonyl oxygen of D243^6.50^. The C=O–Na^+^ distance is 2.8 Å, *versus* 3.6 Å in the TmPPase:IDP:ATC structure. This is mostly because the sodium ion appears to be translated by about ~1.2 Å (see Methods). We believe that these changes are structurally and functionally significant and related to the binding of ATC (see *Asymmetry* below).

### ATC binds in a cleft beside the exit channel

During refinement, we observed extra density near chains A and D (Supplementary Fig. 1c), into which we were able to model ATC. ATC is a rigid multiring system with only two free torsion angles, which we modeled in *trans* geometry, the most prevalent in solution (Supplementary Info. NMR spectra). We could thus, even at this low resolution (Supplementary Fig. 1c), place two molecules of ATC into the (F_o_–F_c_) electron density map. There is some (F_o_–F_c_) density in an equivalent region near chains C and B into which a single ATC could be fitted, but the density there is too poor to fit an ATC dimer, presumably due to molecular disorder in the crystal. We thus focus on ATC**-1** and ATC**-2**, which bind to chains A and D as representing the TmPPase_2_(Mg_5_PP_i_)_2_(ATC_2_)_1_ complex, where the ATC dimer binds asymmetrically to TmPPase.

The cleft binds two ATC molecules stacked tail-to-tail through π-stacking interactions (Fig. 2). Strands β1–2 (loop6–7), loop8–9, and loop12–13 near the exit channel of chain A form a hydrophobic cleft with loops 2–3 and 4–5 on chain D (Fig. 3a). (A similar cleft forms between chains C and B.) In terms of buried surface area; ATC**-1** buries 121 Å^2^ on chain A and 92 Å^2^ on chain D, while ATC**-2** buries 98 Å^2^ on chain A and 94 Å^2^ on chain D (Table 2).

**Figure 3.**
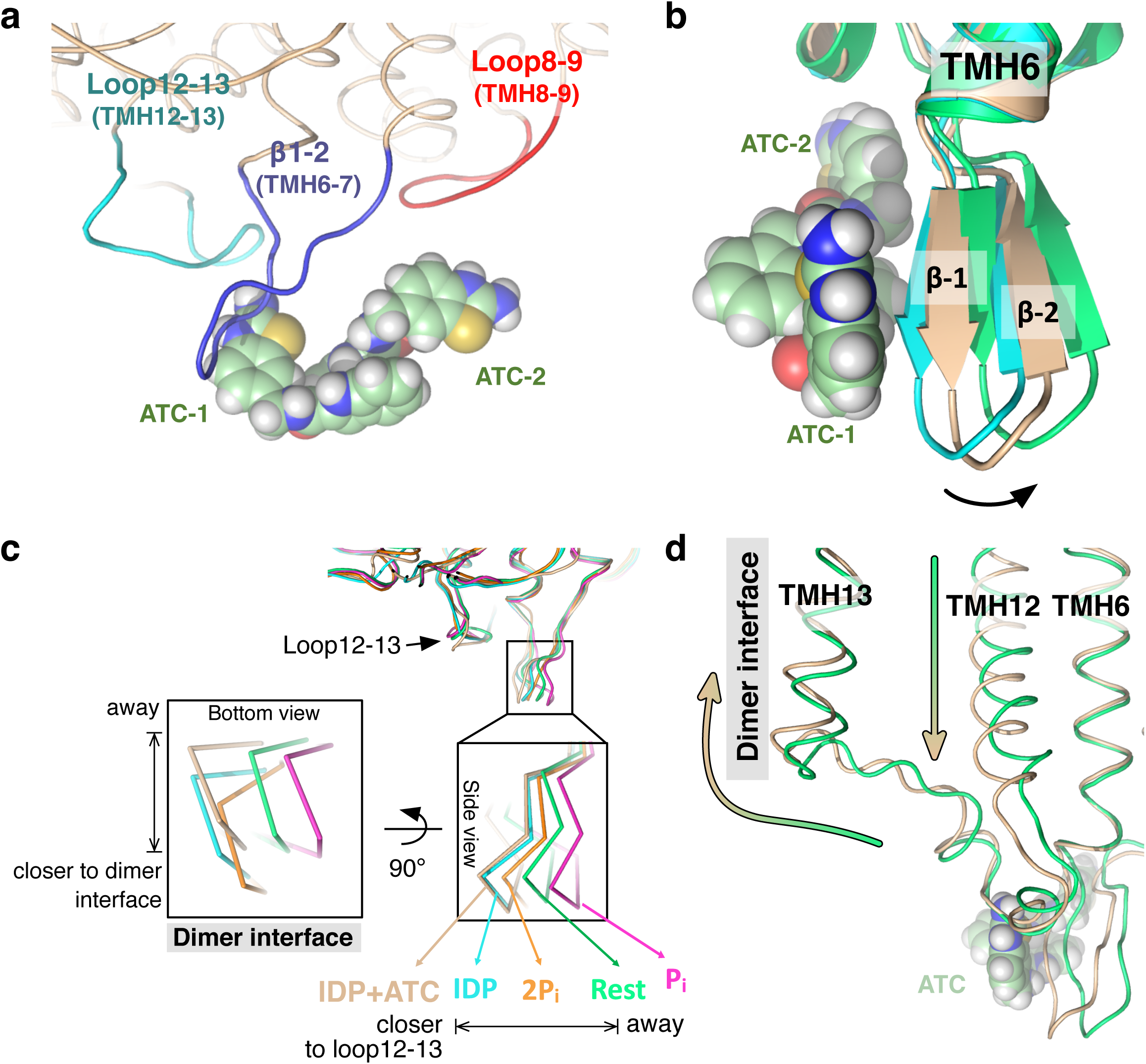
Loop movement caused by the interactions with ATC. (**a**) ATC dimer interaction with β1–2 strand (deep blue), loop8–9 (red), and loop12–13 (cyan). ATC is depicted in space-filling model (pale-green: carbon, blue: nitrogen, red: oxygen and yellow: sulphur). (**b**) Superposition of TmPPase:IDP (cyan) with TmPPase:IDP:ATC structure (wheat) and TmPPase:Ca:Mg (green) showing the large side movement of β1–2 strand (arrow). ATC is depicted as in **a**. (**c**) Superposition of TmPPase structures in all different states showing the closer/away movement of β1–2 strand relative to the loop12-13 and the dimer interface of the protein. Colouring scheme from the left to right: phosphate analogue (WO_4_)-bound state (TmPPase: WO_4_) = pink; resting state (TmPPase: Ca:Mg) = green; two product bound-state (TmPPase:2P_i_) = orange; IDP-bound state (TmPPase:IDP)= cyan; IDP:ATC-bound state (TmPPase:IDP:ATC) = wheat. (**d**) Superposition of TmPPase structures showing the downward movement of TMH12 and away movement of loop12-13 from the resting state (green) to the IDP:ATC-bound form (wheat) of TmPPase. The green-to-wheat coloured arrow shows the movement.

**Table 2.** Major interactions of ATC with chains A and D

| Interaction type | Compound | Chain A |  | Chain D |  |
| --- | --- | --- | --- | --- | --- |
|  |  | Residue | BSA* (Å<sup>2</sup>) | Residue | BSA* (Å<sup>2</sup>) |
| Hydrophobic/<br>Aromatic | ATC-1 | Q268 | 46.3 | M69 | 15.0 |
|  |  | K269 | 34.3 | T73 | 15.9 |
|  |  | I270 | 21.4 | W74 | 15.3 |
|  |  | P530 | 19.1 | N163 | 15.7 |
|  |  |  |  | W164 | 30.1 |
|  | ATC-2 | Q268 | 25.6 | W74 | 45.5 |
|  |  | 277 | 22.2 | Y161 | 32.0 |
|  |  | D351 | 28.6 | F169 | 16.2 |
|  |  | V352 | 21.1 |  |  |
|  | Total BSA* (Å<sup>2</sup>) |  | 218.7 |  | 185.7 |
| Possible<br>hydrogen bond | Compound | Chain A | Distance (Å) | Chain D | Distance (Å) |
|  | ATC-1 (N19) | G528 (C=O) | 3.6 |  |  |
|  | ATC-1 (N20) | G528 (C=O) | 3.8 |  |  |
|  | ATC-2 |  |  | N157 (Oδ1) | 3.6 |
| Salt bridge | ATC-2 (N19) | D351 (Oδ2) | 3.0 |  |  |

\* BSA: Buried surface area: the solvent-accessible surface area (cut-off = 15 $\text{\AA}^2$ ) of the corresponding residue that is buried upon interface formation.

ATC has a 2-aminothiazole group, which is a weak base (p*K*_a_ 4.23 ± 0.03^21^) reported in both neutral and protonated forms in the PDB and CSD. We modeled the 2-aminothiazole groups in the protonated state because both ATCs in the AD interface form ion pairs/hydrogen bonds with chain A (Table 2, Fig. 2b). Chain A D351 (Oδ2) forms a salt bridge with the 2-aminothiazole on ATC-2 while chain A G528 (C=O) forms a bifurcated H-bond to the 2-aminothiazole on ATC**-1** (N19 and N20; see Supplementary Fig. 1d for atom numbering of ATC) (Table 2, Fig. 2b). In chain D, on the other hand, the chief interactions do not seem as strong: a single hydrogen bond is formed, the N157 Oδ1 hydrogen bonds N20 on ATC**-2**, and the remaining interaction are hydrophobic stacking of W164 on ATC**-1**, and the indole of ATC**-2** sandwiched between W74 and W164 and stacking on Y161 (Table 2, Fig. 2c). The interactions are likely due to crystal packing as the binding of ATC to the protein did not change the protein oligomerization in solution (Supplementary Fig. 3). We therefore focus on the interactions of the ATC dimer with chain A.

### Asymmetry

There is recent evidence of functional asymmetry in K^+^-dependent mPPases in the presence of excess potassium^22^. Here we show the first structural evidence of asymmetry. The IDP-bound state has the most symmetrical structure (the lowest rmsd values) between each monomer compared to others (Table 3, Supplementary Fig. 4a). However, the binding of ATC makes the dimeric protein asymmetric, especially in the loops that interact with the inhibitor (loops8–9, β1–2 and 12–13). The maximum rmsd is 1.6 Å (Table 3). Structural alignment between monomers of other states (the resting state, 2P_i_-bound state and WO_4_^-^ bound state) of those loops also shows more structural symmetry. In addition, there is a change in the binding mode of the Na^+^ at the ionic gate (see above).

**Table 3.** Root mean square deviation (rmsd) of chain A and B of loops in different TmPPase structures.

| Loop | Rmsd ($\text{\AA}$)<sup>1</sup> | | | | |
| --- | --- | --- | --- | --- | --- |
|  | TmPPase:IDP:ATC<br>(AB) | 5lzQ<br>(AB) | 5lzR<br>(AB) | 4AV3<br>(AB) | 4av6<br>(AB) |
| 2-3 (71-75) | 0.382 | 0.184 | 0.444 | 0.331 | 0.261 |
| 4-5 (153-172) | 0.905 | 0.242 | 0.579 | 0.369 | 0.322 |
| β1-2 (6-7) (266-276) | 1.630 | 0.486 | 1.237 | 0.613 | 0.503 |
| 8-9 (342-359) | 0.743 | 0.405 | 0.695 | 0.337 | not applicable<sup>2</sup> |
| 12-13 (520-540) | 0.442 | 0.367 | 0.406 | 0.455 | 0.364 |

<sup>1</sup> Comparisons were done by superimposing helix 7 (281-311), which moves least between the structures
<sup>2</sup> Missing loop from residue 349-357

ATC binds as a dimer in chain A, holding the β1–2 loop (Q268, K269, I270, and Q277), loop8–9 (Q348, D351, and V352), and loop12–13 (G528, P530, and P531) together (Fig. 3a; Supplementary Fig. 4b, c). Structural comparison of the loops that interact with ATC with the equivalent loops in TmPPase:IDP showed that the β1–2 and 12–13 loops in chain A move the most, with an RSMD per Cα of 1.15 and 0.80 Å respectively (Supplementary Table 2). This is due to the interactions with the ATC dimer (Fig. 3b, Table 2). In contrast, the RMSDs/Cα for the other loops that interact with ATC are smaller (Table 3). When all the TmPPase states are compared, it is clear that the β1–2 loop moves closer/away relative to loop12–13 and the dimer interface of the protein (Fig. 3c). In particular, the binding of ATC causes the β1–2 strand of chain A to move away from the dimer interface relative to its position in TmPPase:IDP and TmPPase:P_i2_ to roughly the same angle as in the resting and P_i_ structures (Fig. 3c, inset bottom view; periplasmic side) – when the exit channel must be closed. The movement of chain A β1–2 due to the binding of the ATC dimer thus create a hydrophobic clamp that locks the TmPPase exit channel in a closed state after substrate binding. In contrast, it is reasonable to assume that ATC does not bind to chain B because the β1–2 strand is moving away from the loop12–13, closer to its position in the resting state (Table 3 and Supplementary Fig. 5a). This movement appears to prevent ATC binding because it leads to the protrusion of Q268 and Q277 that occlude the ATC binding site (Supplementary Fig. 5b).

### Pharmacology of ATC and analogues at TmPPase

Together with ATC, we identified four of its analogues (compound **2**, compound **3**, compound **4**, compound **5** (Fig. 4a). A set of four analogues (**6**, **7**, **8**, and **9**), three of them brominated, were then designed for structural studies. Their IC_50_s were at least 20 times worse than the original hit ATC (Fig. 4b, c), and they did not help us solve the structure. They do, however, provide some insight into the structure-activity relationship (SAR) for this family of compounds on TmPPase.

**Figure 4.**
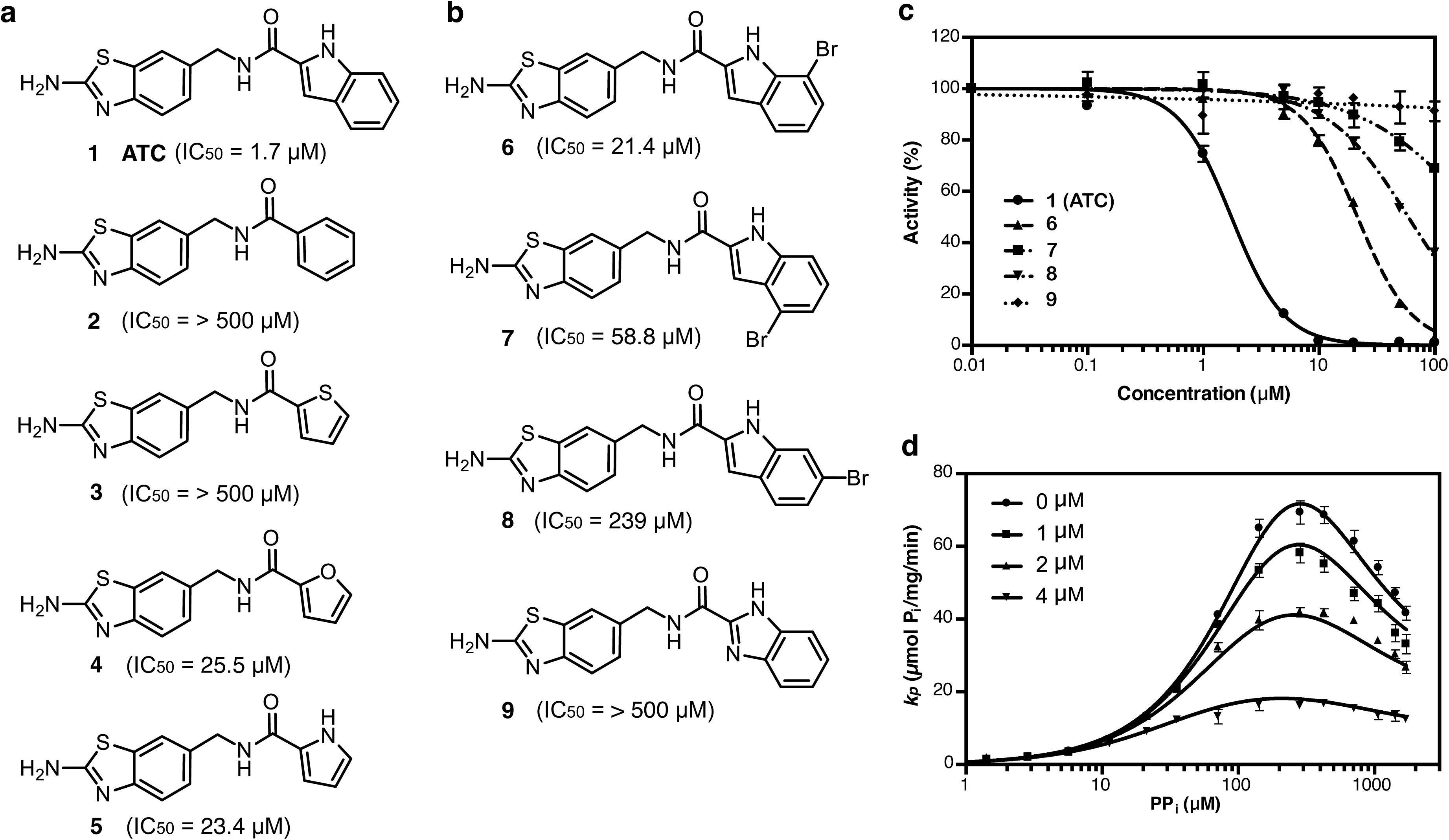
Compound library and inhibition activity against TmPPase. (**a**) Screening hits. (**b**) Analogues developed for structural studies. (**c**) Inhibition curve of ATC, compounds **6–9**. (**d**) Kinetic inhibition plot for the inactivation of TmPPase at four different concentrations of ATC. The solid lines represent nonlinear regression fits with the residuals standard deviation (Sy.x) of 2.0. All data shown as mean ± SD with n=3 replicates.

The suggested binding modes also explain the structure-activity relationships (SARs) of the compounds presented in Figure 4. First, they highlight that the hydrogen bonding functionality of the indole ring is important; compounds **2** and **3** that lack such a functionality are inactive. In the TmPPase:IDP:ATC structure, there are two asparagine residues (N163 and Q268) near the indole nitrogen of ATC, which would explain not only the lack of activity of compounds **2** and **3**, but also the tolerance of both hydrogen bond donor (nitrogen; compound **5**) and acceptor (oxygen; compound **4**) functionalities in this position. Second, the aromatic nature of the indole ring seems important for activity: compounds **4** and **5** that include suitable hydrogen bonding functional groups but not a bulky ring structure are approximately 10-fold less active than ATC. The indole rings of ATC**-1** and ATC**-2** form π-π stacking interactions with each other, and removing the benzene ring from the structure weakens this interaction. Third, the compounds **6**, **7** and **8** with bromine substitutions are from 10-fold to 100-fold weaker binders than ATC, suggesting the importance of the unsubstituted indole ring. The bromine substitution might weaken the π–π stacking interactions observed in the structure by altering the shape and location of the π-electron cloud, but also directly clash with the loops 2–3 and 4–5; the latter is especially probable with the weakest brominated compound **8**. Finally, benzimidazole substitution instead of an indole yields fully inactive compound **9**. This might again be due to the loss of the observed stacking of the indole rings of ATC**-1** and ATC**-2**: the benzimidazole group could be both protonated at physiological pH, leading to repulsive interactions.

### ATC binds tightly to the E_2_S form

As ATC is a potent inhibitor, we further characterized its effect on the rate of substrate (PP_i_) hydrolysis using a range of compound concentrations (0.0 – 4.0 μM). We performed the kinetic assay using PP_i_ concentration of 0 – 1714 μM at 71°C with 2 minutes of reaction time (Fig. 4d). The simultaneous analysis of the data for all inhibitor and substrate concentrations is consistent with a model in scheme 1 with equation (1) and n=2 (Table 4), with the exception that K_I1_ is not fit (when we included it, the residuals did not improve, and its mean value is >0.5 mM; data not shown). The standard deviation of the residuals (Sy.x) for the dimer fit is 2.0 for scheme 1 where n=2, but 4.3 when n=1, and the curves clearly do not fit (data not shown). This means that ATC binds as a dimer to TmPPase. ATC clearly binds tightest to the E_2_S complex (*K*_I2_ = 1.8 ± 0.3 μM) and the binding to E_2_S_2_ is 5-fold weaker (10.0 ± 1.1 μM) and no binding to E_2_ (Table 4). ATC is thus an uncompetitive dimeric inhibitor with a clear preference for binding to the single-substrate bound form. After the binding of substrate to monomer A (E_2_S), monomer B of TmPPase showed higher affinity to the substrate to form E_2_S_2_ complex as the α value is very low (0.12). However, after both monomers bind substrate (E_2_S_2_), the hydrolysis rate decreases by twenty-fold (β = 0.05) compared to the rate of hydrolysis of the E_2_S complex (*k*_p_ = 239.2 ± 32.2 μmol P_i_/mg/min) (Table 4).

**Table 4.**
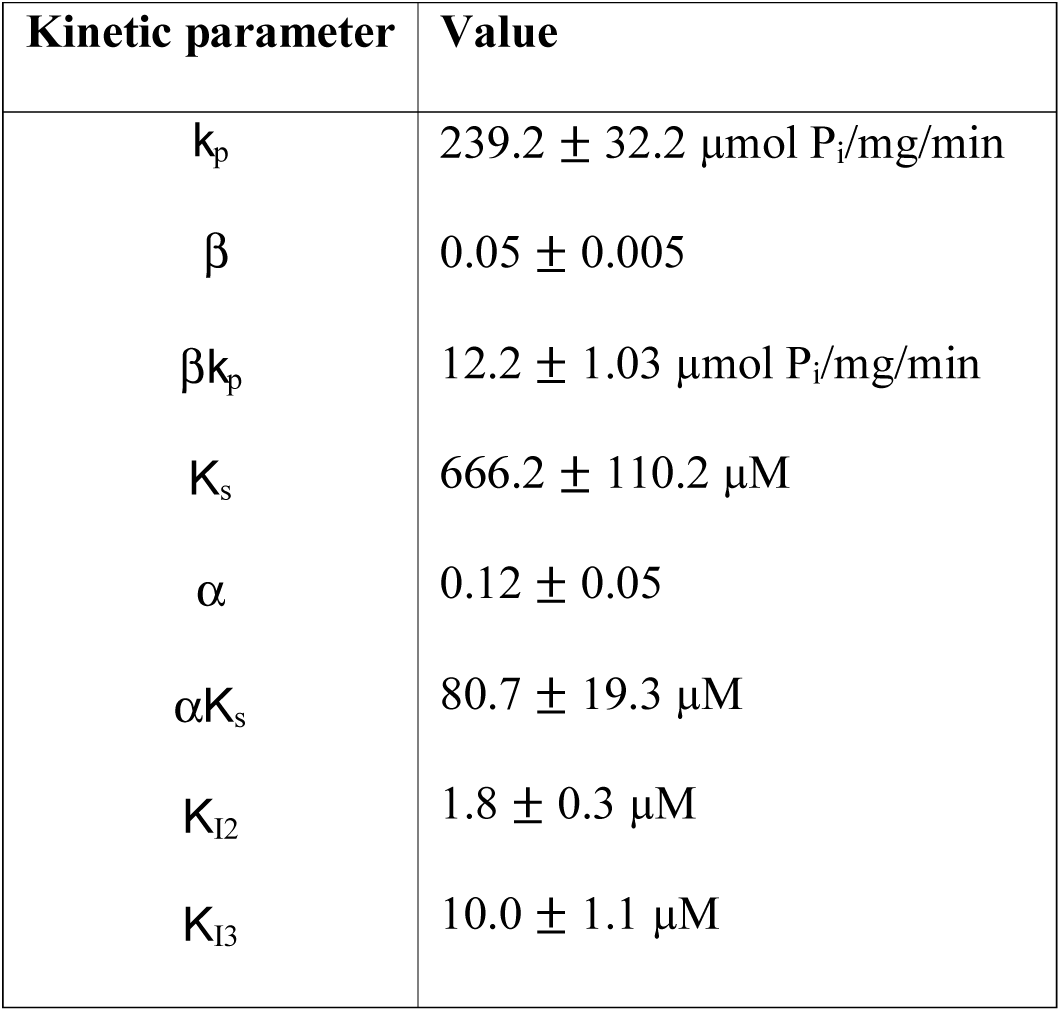
Kinetic parameters of TmPPase activity and ATC inhibition constants

Overall, it is clear that TmPPase shows positive cooperativity for substrate binding but negative cooperativity for hydrolysis, one where only the E_2_S form binds the ATC inhibitor tightly. In the crystal structure we solved, ATC is bound to the E_2_S_2_ form, not to the E_2_S form. This is probably because the concentration of ATC added for crystallization was higher (1 mM) than the binding affinity of ATC to the E_2_S_2_ form (10.0 μM) or because the ATC binding to E_2_S form can still induce the conformational change in the other monomer that allows further substrate binding (see Discussion). This is consistent with the fact that we were unable to crystallise the TmPPase:ATC complex except in the presence of IDP as a competitive inhibitor. ATC is thus the first non-phosphorus and the first allosteric inhibitor identified for mPPase.

### Use of ATC in designing inhibitors of parasitic mPPases

ATC would be a potential lead as an antiparasitic agent if it were active against protozoan parasites like *Plasmodium* spp. Preliminary studies with isolated *P. falciparum* membranes showed that ATC had an IC_50_ around 80 μM (Vidilaseris, *unpubl.*), *i.e* at least 40-fold lower than in our test system *T. maritima* (Fig. 4a). In the *P. falciparum* survival assays, ATC did not show any anti-plasmodial activity (Supplementary Fig. 6). On the other hand, knockout of mPPases by RNAi in *T. brucei*^23^ and in *P. falciparum* (Totanes, *unpubl.*) demonstrate that functional mPPase is required for survival and virulence.

To explain these findings and study the potential specificity determinants at the binding sites of ATC**-1** and ATC**-2** across mPPases, we retrieved the sequences of 16 mPPases from protozoan parasites (Supplementary Fig. 7). Within these sequences, the areas corresponding to the loops 2–3, 4–5, 6–7 (i.e. β1–2 strand), 8–9 and 12–13 showed a high degree of variation (Fig. 5a and Supplementary Fig. 7). As the binding site of ATC**-1** and ATC**-2** is located on these loops (6–7, 8–9 and 12–13 of chain A), it is unlikely that ATC would bind to the mPPases of pathogenic parasites as it does in the current TmPPase structure. To validate this, we built a homology model of *P. falciparum* mPPase (PfPPase; Fig. 5b). The loops 4–5, 6–7 and 12–13 of the modelled structure are notably shorter than those of TmPPase: the ATC binding site is not present, as loop 6–7 contributes about 2/3 of the interface (Table 2). This not only explains why ATC does not inhibit PfPPase, but also supports the notion it functions asymmetrically, preventing a full catalytic cycle from occurring by binding to one chain and preventing the exit channel from opening.

**Figure 5.**
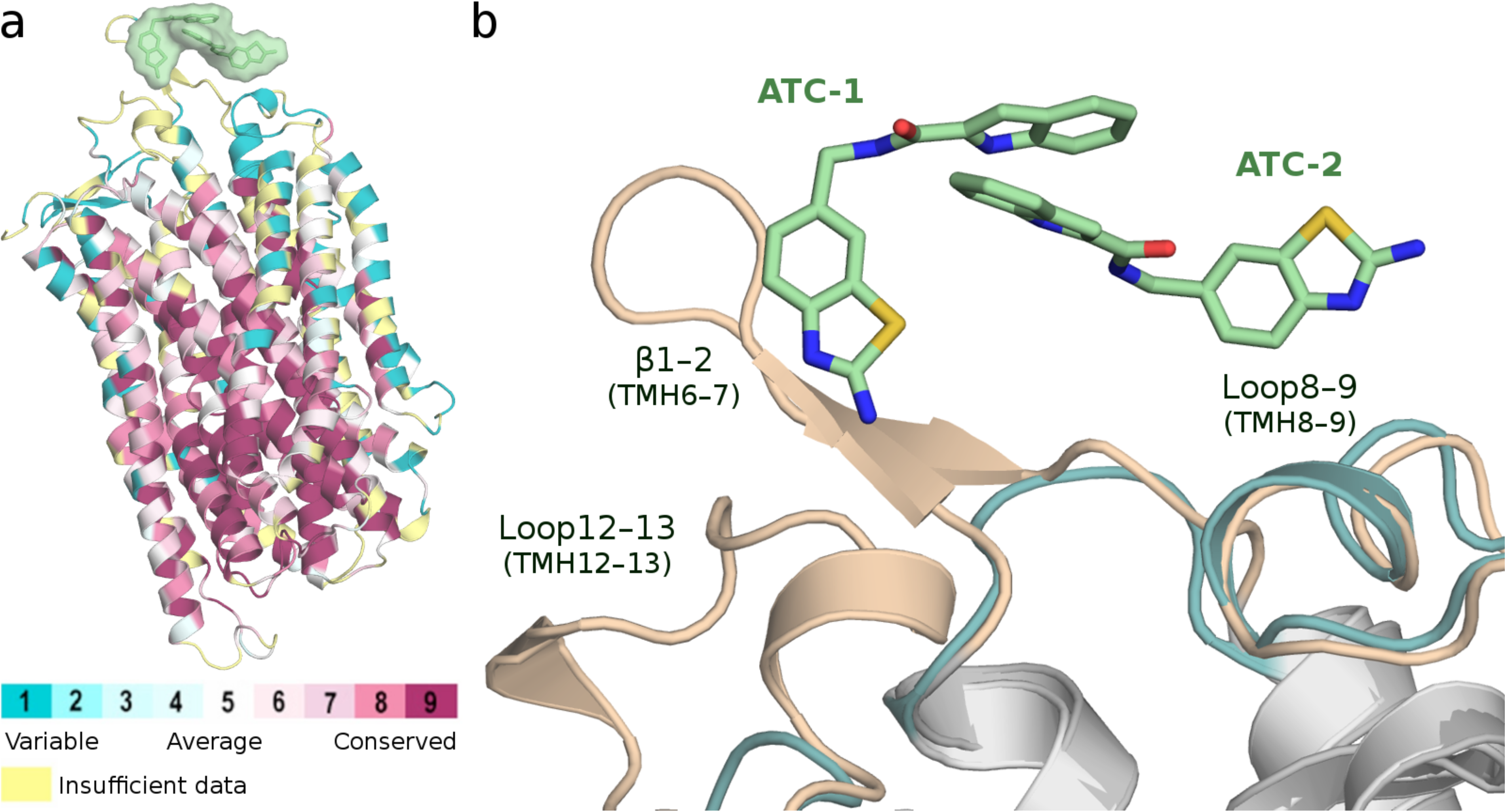
The conservation of ATC binding site among mPPases. (**a**) The structure of TmPPase coloured according to the sequence conservation among the set of 18 pathogenic mPPases. The pale-green surface indicates the location of the ATC binding site. (**b**) Comparison of the ATC binding sites of the crystal structure of TmPPase (wheat) and a homology model of PfPPase (teal). Carbon atoms of ATC are presented as pale-green sticks. Blue: nitrogen, red: oxygen and yellow: sulphur. View as in Figure 2A.

## Discussion

### The first novel inhibitors of integral membrane pyrophosphatases

mPPases are potential targets for anti-protozoan drug design as they do not occur in humans and other mammals, are essential in *P. falciparum* (Totanes, *unpubl.*) and render *T. gondii* non-infectious^19^ Bisphosphonate derivatives (PP_i_ analogues) can inhibit mung bean mPPase^24^ and prevent the proliferation of *T. gondii*^25^ and other protozoan parasites^26,27^. However, they also inhibit human enzymes such as farnesyl pyrophosphate synthase, so they are not viable as anti-parasitic drug candidates (for a review, see ^13^). This study reports the first non-phosphorus inhibitors of mPPase, which were identified through a screening process. The best compound (ATC), binds and inhibits both the E_2_S and E_2_S_2_ complexes, with inhibition constants *K*_i2_ and *K*_i3_ of 1.8 and 10.0 μM, respectively (Table 4). The overall IC_50_ is 1.7 μM. It appears to act in an allosteric manner. This mode of inhibition has been observed for an inhibitor of soluble PPase from *Mycobacterium tuberculosis* (MtsPPase)^28^, but to our knowledge, this is the first one observed in mPPase.

### Identification of alternate inhibitory mechanisms and evidence for allostery in mPPases

ATC clearly binds beside the exit channel (Fig. 2 and Fig. 3). Similar types of inhibition occur in other membrane proteins. For instance, the voltage-dependent K_v_11.1 (hERG) potassium channel that mediates the repolarizing current in the cardiac action potential is easily inhibited by a wide variety of drugs, leading to drug-induced long QT symptom and lethal arrhythmia. The molecules, such as quinolone and macrolide antibiotics, bind to hydrophobic pockets on the side of the selectivity filter^29,30^. Recently, a structural study found inhibitors that targeted a hitherto-unknown allosteric transmembrane site in MsbA, an ATP-binding cassette (ABC) transporter. They lock MsbA in an inward facing conformation that prevents transition to the outward-facing conformation during lipopolysaccharide transport^31^. The inhibitor is wedged in the membrane-exposed pocket formed between TMH4, TMH5, and TMH6 of each subunit. This binding uncoupled the nucleotide-binding domains (NBDs), which adopt an asymmetrical conformation.

Similar interactions to the tail-to-tail π-stacked ATC dimer that binds to TmPPase have been observed in soluble protein-inhibitor complexes as well. For example, Stornaiuolo, Kloe^32^ found an isoquinoline derivative that formed a π-stack of three identical molecules that bound to one binding site of acetylcholine-binding protein. A quinazoline derivative formed a π-stack of five molecules to the lysine-specific demethylase 1 (LSD1) — RE1-silencing transcription factor co-repressor (CoRESt) complex^33^, and a phenylpyrrole derivative forms a head-to-tail π-stacked dimer that binds in the two-fold cavity in soluble hexameric *M. tuberculosis* PPase. Similarly to the TmPPase complex, this inhibitor appears to act uncompetitively^28^, but the interaction clearly has 1:1 stoichiometry, with the binding site is 20 Å from the nearest pyrophosphate^28^. The asymmetric binding of the ATC dimer to one monomer of TmPPase is similar in general.

A recent study has shown that all mesophilic mPPases show substrate inhibition with increased *K*_m_ and decreased *V* when two molecules of substrate are bound in the presence of 50 mM K^+ 22^. The binding of S to E_2_S is weaker than to E_2_, and *k*_cat_ for E_2_S_2_ is lower than for E_2_S. They interpreted this to mean that substrate binding to one monomer causes a conformational change in the other that inhibits its interaction with the substrate. Our ATC-bound structure (Fig. 2 and Fig. 3) and inhibition data (Fig. 4 and Table 4) are similar: an ATC *dimer* binds only to one monomer in the TmPPase_2_:(Mg_5_PPi)_2_ dimer. The second site is unavailable in the substrate-bound form, and neither monomer binds ATC before substrate binds. However, our data differ from theirs in that binding of the first substrate molecule appears to be much weaker than binding of the second as α is 0.125, while the hydrolysis rate in the E_2_S_2_ complex is twenty-fold slower (Table 4).

How does ATC exert its effects? After substrate binds, its binding to chain A *via* loops 6–7, 8–9 and 12–13 creates a hydrophobic clamp that locks the exit channel in the closed state and TMH12 in the down state. We posit that the mechanistic implications of these events are as follows. β1 extends from TMH6, which contains D243^6.50^, D246^6.53^, and D247^6.54^, all of which coordinate Na^+^ in the gate. In the ATC-bound structure, Na^+^ is coordinated by four oxygens from three Asp side chains and a serine, not five as in the TmPPase:IDP structure. Furthermore, in our current structure, D246^6.53^, D703^16.46^, and Na^+^ move up to 1.2 Å from their positions in the TmPPase:IDP structure (Fig. 1c). This corresponds to a new state that, we believe, keeps the exit channel closed. Our current model^10^ posits that, during ion pumping, TMH12 moves downward by at least 2 Å^9^. Comparing the relative positions of the ATC-bound and resting states suggests that this movement of TMH12 makes loop12-13 move towards the dimeric interface of the protein and TMH13 move “up” towards the cytoplasmic side (Fig. 3d). This would induce equal conformational changes in monomer B in the TmPPase:IDP. However, ATC binding locks monomer A TMH12 in the “down” state. Because of its “down” position, loop12–13 and TMH13 can still trigger the conformational changes of monomer B into a state that binds substrate more tightly. However, further catalysis is impossible because monomer B cannot undergo a full catalytic cycle: substrate can bind, but the full motions required for catalysis are impossible. Diagnostic of these motions is the conformation that leads to binding of ATC. In agreement with the structure, the kinetic data shows that ATC dimer only binds to one of the monomers, and binds most tightly to the TmPPase_2_(Mg_5_PP_i_) state (Table 4).

The requirement for potassium for maximal activity and for substrate inhibition is also explained by these structures. The amino acid responsible for K^+^ dependency (A495^12.46^ in TmPPase) is located in TMH12, which is directly linked to the dimer interface (loop12-13 and TMH13). We suggest that, in the presence of K^+^, the downward movement of TMH12 upon substrate binding and ion pumping induces conformational changes in the monomer-monomer interface by the “away” movement of loop12-13 and upward movement of TMH13 (Fig. 4d). This is then propagated to other TMHs. In the absence of K^+^, such coupling does not occur. These changes increase affinity and activity. They also provide a possible structural explanation for the Na^+^/H^+^ pumping enzymes^22,34^, where H^+^-pumping is not inhibited by 100 mM Na^+^ and 50 mM K^+^. One way this might happen is through a conformation similar to that observed here: the binding of PP_i_ to monomer A and pumping of, say, a proton would drive changes in the loop12-13 and TMH 13 such that the conformation of monomer B became suitable for binding PPi and pumping Na^+^.

In conclusion, we propose the following model for TmPPase catalysis and its inhibition by ATC (Fig. 6). Substrate binding to monomer A leads to pumping in monomer A and induces a conformational change in monomer B that increases its affinity for PP_i_. After PP_i_ binds in monomer B, PP_i_ hydrolysis occurs in monomer A and ion pumping in monomer B. The product of catalysis (2P_i_) is released from monomer A, allowing hydrolysis and subsequent product release in monomer B. The ATC dimer can only bind when PP_i_ is bound to monomer A. Nonetheless, the A:PP_i_ form can still induce the conformational change in monomer B that allows PP_i_ binding. However, since the ATC binding locks monomer A in the down state, no further catalysis can occur.

**Figure 6.**
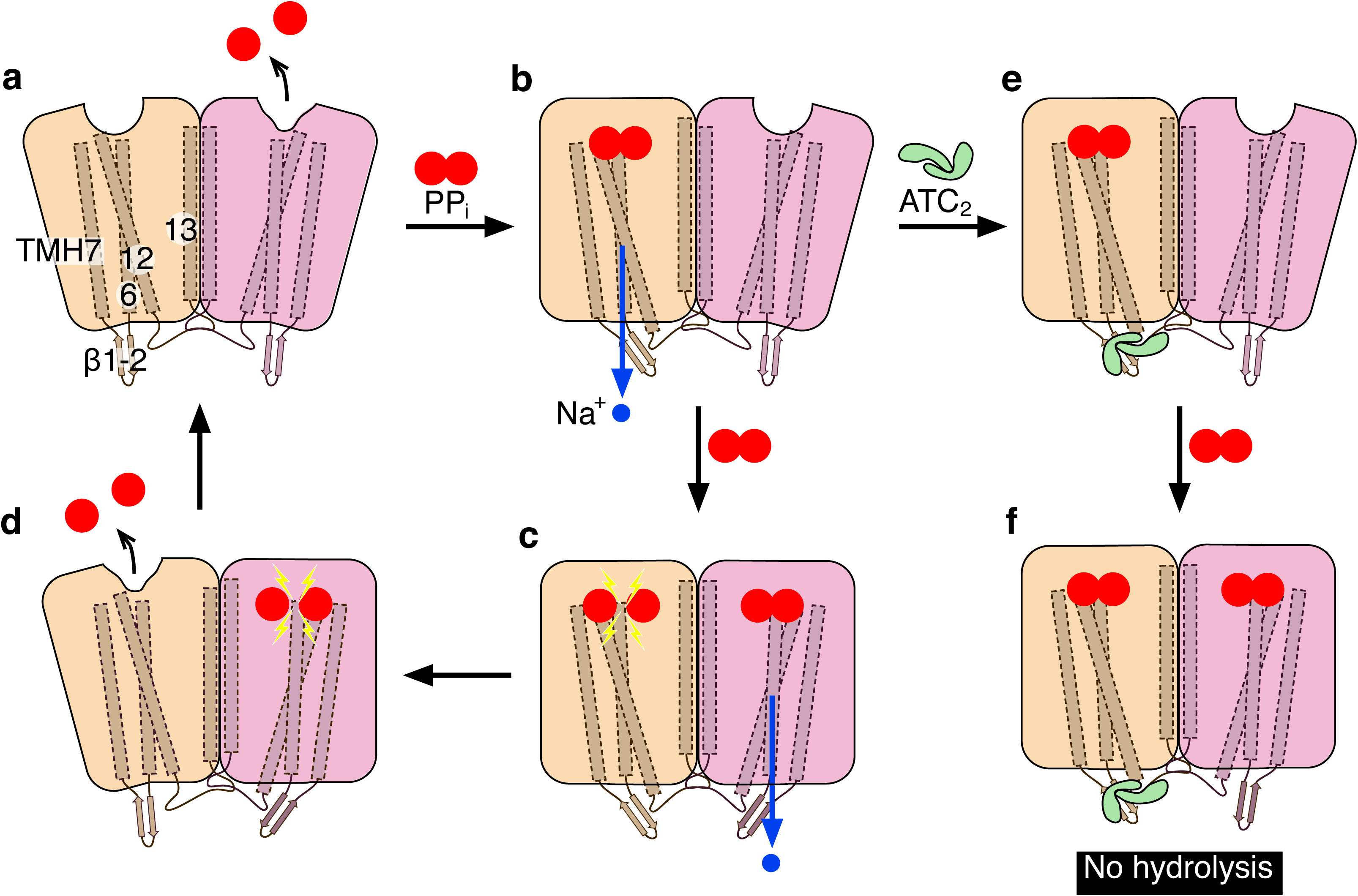
Catalytic scheme of TmPPase and its inhibition by ATC. The binding of substrate (PP_i_; two joined red circles) to monomer A (wheat colour) (**a**) leads to Na^+^ pumping (blue circle and blue arrow for the pumping direction) and induces a conformational change in monomer B (pink colour) (**b**) which increase its affinity for substrate binding. The Na^+^ pumping causes the substrate hydrolysis (yellow lightning symbol for this event) in monomer A, while the substrate binding in monomer B leads to its Na^+^ pumping (**c**). After hydrolysis, the monomer A released its hydrolysis products (2P_i_; red circles) while monomer B proceed to the hydrolysis event (**d**) and then moves back to the state in a. The ATC dimer binds to its binding site after substrate binding and Na^+^ pumping in monomer A (**e**) which lock this monomer in the “down” state. Even though no hydrolysis occurred in monomer A due to the ATC binding, its conformational change still able to induces a conformational change in monomer B that increase its substrate affinity (**f**). The numbering with white background corresponds to transmembrane helix number.

Our novel structure of TmPPase with IDP and ATC has opened up new routes to drug discovery by serendipitously demonstrating that there are alternative ways to inhibit mPPase. Although the ATC-binding interface is not preserved (Fig. 5), our data nevertheless indicate that binding at the exit channel can inhibit mPPases. The work shows how extracellular loop binding regions can have significant effects on intracellular substrate binding and *vice versa*. Finally, by stabilizing asymmetric complexes, ATC provides a route into determining other steps along the reaction pathway and thus deriving a full catalytic model – one that may unify the Na^+^/H^+^ and Na^+^ pumpers as outlined above.

## Methods

### Discovery of ATC and its analogues

The screening process that led to the discovery of compound **1** (ATC) will be described in more detail elsewhere. Briefly, we tested compounds using a recently developed 96-well plate assay with TmPPase as a model system^35^. The screening collections included compounds synthesized in our laboratories. The identification of ATC as an inhibitor was followed by the test of analogues **2–5**. ATC and its analogues were initially developed for their antimicrobial, antiviral and anticancer activities as reported elsewhere^36^.

### Compound synthesis

The purity of all the compounds tested was above 95%. The synthesis of ATC (compound **1**) and compounds **2–5** has been described elsewhere^36^. The synthesis of compounds **6–9** is described in supplemental material. For synthesis, all reactions were carried out using commercially available starting materials (Sigma-Aldrich, Schnelldorf, Germany; BioFine International Inc, Vancouver, Canada, Fluorochem, Hedfield, UK) and solvents without further purification. Column chromatography was performed with an automated Biotage high performance flash chromatography Isolera One (Uppsala, Sweden) using a 0.1-mm path length flow cell UV-detector/recorder module (fixed wavelength 254 nm). Analytical thin layer chromatography (TLC) was carried out using 0.2mm silica gel plates (silica gel 60, F_254_, Merck KGaA, Darmstadt, Germany). Nuclear magnetic resonance spectra (^1^H NMR and ^13^C NMR) were recorded on a Bruker Ascend 400 (Bruker Corporation, Billerica, Massachusetts, USA). ^1^H NMR at 400 MHz and ^13^C NMR at 100 MHz. High resolution mass spectra (HRMS) were measured on a Waters Synapt G2 (Waters Corporation, Milford, Massachusetts, USA) and reported for the molecular ions [M+H]^+^.

### Protein expression and purification

Expression and purification of TmPPase are described elsewhere^37,38^. In brief, His-tagged TmPPase in pRS1024 plasmid under the control of the PMA1-promoter was freshly transformed into *Saccharomyces cerevisiae* strain BJ1991. The cells were grown overnight in 250 ml of selective synthetic complete drop-out (SCD) media and then added to 740 ml of 1.5× YP media containing 2% glucose. The cells were further grown at 30 °C for 8h, collected by centrifugation (4,000 rpm, 10 min) and lysed at 4 °C using a bead beater with 0.2 mm glass beads. The membrane fraction was collected by ultracentrifugation (100,000 × *g*, 45 min) and the pellets were resuspended in buffer containing 50 mM MES-NaOH pH 6.5, 20% (v/v) glycerol, 50 mM KCl, 5.2 mM MgCl_2_, 1.33 mM dithiothreitol (DTT), 2 μg ml^-1^ (w/v) pepstatin-A (Sigma) and 0.334 mM PMSF (Sigma). The membranes were solubilized using the ‘hot-solve’ method^38^ at 75 °C for 1.5 h in solubilisation buffer (50 mM MES-NaOH pH 6.5, 20% (v/v) glycerol, 5.33% (w/v) *n*-dodecyl-β-D-maltopyranoside (DDM) (Anatrace)). After centrifugation to remove denatured proteins, KCl (to a final concentration of 0.3 M) and 2 ml of Ni-NTA beads (Qiagen) were added and incubated at 40 °C for 1.5 h to each 40 ml of the solubilized protein, and then loaded into an Econo-Pac^®^ column (Bio Rad). Then the column was washed with 2 × CV (column volume) of washing buffer (50 mM MES-NaOH pH 6.5, 20% (v/v) glycerol, 50 mM KCl, 20 mM imidazole pH 6.5, 5 mM MgCl_2_, 1 mM DTT, 2 mg ml^-1^ (w/v) pepstatin-A, 0.2 mM PMSF and 0.05% DDM (Anatrace) and eluted with 2 × CV of (50 mM MES-NaOH pH 6.5, 3.5% (v/v) glycerol, 50 mM KCl, 400 mM imidazole pH 6.5, 5 mM MgCl_2_, 1 mM DTT, 2 mg ml^-1^ (w/v) pepstatin-A, 0.2 mM PMSF and 0.5% octyl glucose neopentyl glycol (OGNPG, Anatrace), 1 mM DTT, 2 mg ml^-1^ (w/v) pepstatin-A, 0.2 mM PMSF and 0.05% DDM (Anatrace).

### Activity measurement and kinetic analysis

TmPPase activity, compound screening and kinetic experiments were done using the molybdenum blue reaction method in a purified protein solubilised in DDM^35^. Initial reaction rates of TmPPase were determined using PP_i_ as substrate and ATC as inhibitor at varying concentrations. The concentration of MgCl_2_ and Na_4_PP_i_ required to maintain 5 mM free Mg^2+^ at pH 8.0 were approximated as described previously^39^. The reaction was done in reaction buffer (60 mM Tris-HCl pH 8.0, 5 mM free Mg^2+^, 100 mM KCl, 10 mM NaCl) at 71 °C for 2 min.

For analysis, we used a variant of Artukka, Luoto^22^, which assumed different kinetic behavior of each monomer active sites of the mPPase dimer, with the addition of inhibitor (Scheme 1). E_2_ represents the dimeric enzyme, S represents substrate (Na_4_PP_i_), I represents inhibitor (ATC), n represent the amount of ATC binds to the enzyme, *K*s is a microscopic Michaelis—Menten constant, and *k_p_* is per-site maximum rates for the substrate complex. Alpha (α) is the factor relating the binding of the first substrate molecule (*K*S) to the binding of the second, and β the factor relating the rate of hydrolysis of the E_2_S complex (*k*_p_) to that of the E_2_S_2_ complex, and *K*_I1_, *K*_I2_, *K*_I3_ the inhibition constants for binding to the E_2_, E_2_S and E_2_S_2_ complexes.

**Scheme 1.**
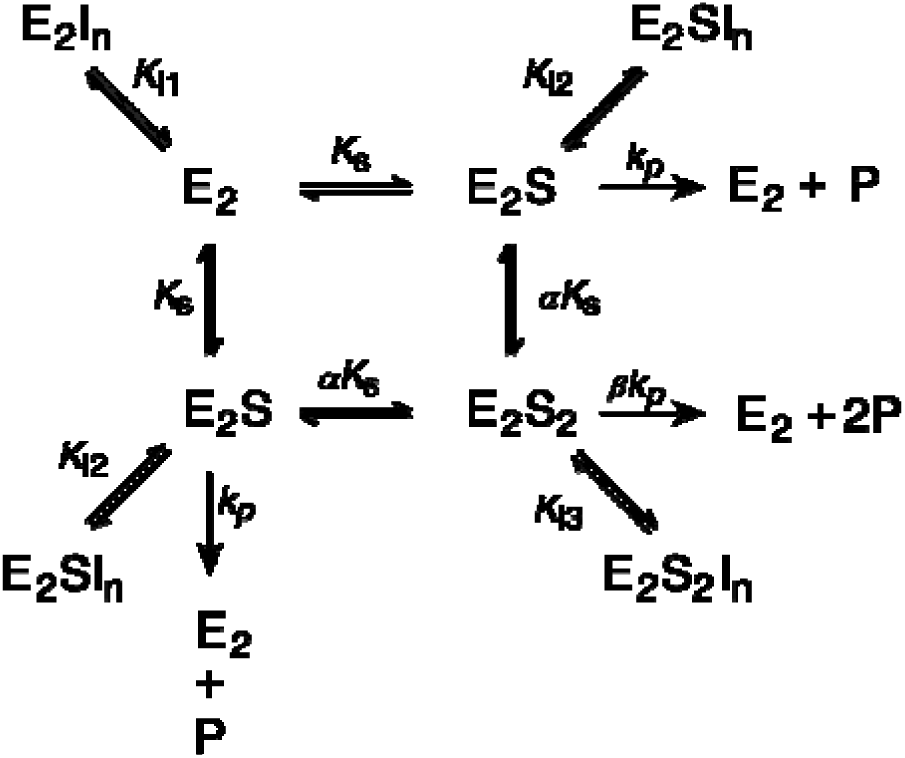
Substrate binding, hydrolysis and inhibition at two active sites of a dimeric TmPPase.

For the mechanism in scheme 1 with n=2 (*i.e.* only inhibitor dimers bind), the rate equation (1)^40^ is:

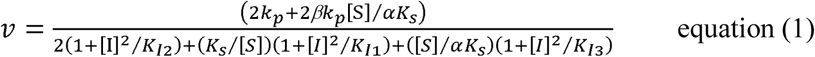

where [S] and [I] correspond to substrate and inhibitor concentrations, respectively and the equilibrium and Michaelis-type constants are as in scheme 1. The kinetic data were fit together to obtain all parameter values and their standard errors by nonlinear regression as a function of inhibitor (0, 1.0, 2.0, and 4.0 μM) and substrate (0 — 1714 μM) concentrations using Prism 6.0 (GraphPad software).

### Crystallization, structure determination and analysis

For crystallization, the purified protein was buffer exchanged to the crystallization buffer (50 mM MES-NaOH pH 6.5, 3.5% (v/v) glycerol, 50 mM KCl, 5 mM MgCl_2_, 2 mM DTT and 0.5% OGNPG) on a Micro Bio-Spin 6 column (Bio-Rad) and then diluted to a concentration of 10 mg ml^-1^. Prior to crystallization, 1 mM imidodiphosphate (IDP) and 1 mM ATC were added, the solution incubated on ice for 10 min, and centrifuged for 20 min (16,000 × *g*, 4 °C). Crystallization trials were done at 22 °C by sitting drop vapour-diffusion method using commercial screens, MemGold^TM^ and PGA screen^TM^ (Molecular Dimensions) in MRC 2-well crystallisation plates (Swissci) with a Mosquito robot (TPP Labtech) and the drops were monitored using the minstrel DT UV imaging system (Formulatrix). Crystal hits appeared in the MemGold^TM^ screen under different conditions. Crystallization conditions were then optimised to get better crystals by varying pH, buffer, salt and PEG concentrations using the vapour-diffusion method in 1 μl + 1 μl (protein-mother liquor) drops in a 24-well plate at room temperature. Harvestable crystals appeared within a week and were frozen after soaking in cryo-protectant (mother liquor + 20% glycerol) or directly from mother liquor. The best diffracting crystal appeared from a solution containing 0.1 M MES pH 6.5, 0.1 M NaCl, 33% polyethylene glycol (PEG) 400, and 4% ethylene glycol.

X-ray diffraction data were collected at ESRF Grenoble (France) on the ID29 beamline at 100 K on a Pilatus 6M detector. 1400 images were collected at an oscillation angle of 0.1°. The data were merged and scaled using X-ray Detector Software (XDS)^41^ and the structure solved by molecular replacement with Phaser^42^ using the IDP-bound state of TmPPase structure (5LZQ)^10^ as the search model. Phaser found a unique solution in space group P2_1_2_1_2_1_ with four monomers in the asymmetric unit forming two dimers. The structure was built and refined using phenix. refine^43^ and Coot^44^. The final model has R_work_/R_free_ of 22.5%/26.8% with 94.1% of residues in the favoured and 5.7% in the allowed region of the Ramachandran plot (Table 1).

The structural interface and assembly of the protein were analysed using the PDBe PISA server (http://www.ebi.ac.uk/pdbe/pisa/)^45^. Internal pocket analysis was done using the Splitpocket server (http://pocket.med.wayne.edu/patch/)^46,47^. Structural alignments were done in PyMOL with the default setting using the monomer structure, except if stated differently. All structural figures were produced using PyMOL (https://pymol.org/2/).

### Comparison with pathogenic mPPases

The sequences of TmPPase and 16 pathogenic mPPases were aligned using Clustal Omega software^48^, and conservation analysis was conducted and visualized with the ConSurf server^49,50^ with default parameters. The initial pairwise sequence alignment of TmPPase and *P. falciparum* mPPase (VP1; PfPPase; sequence identity 42%) was produced with Discovery Studio 4.5 software^51^using the Align123 algorithm. Homology modelling was carried out with MODELLER 9v14^52^ with default settings in the Discovery Studio platform. The pairwise alignment was revised during the course of model building; in total nine models were constructed in three modelling rounds. The Modeller DOPE scores did not differ significantly among these models, and a representative model was selected to minimize strain in the loops connecting transmembrane helices.

## Acknowledgements

This work was supported by the grants from the Jane and Aatos Erkko Foundation and the BBSRC (BB/M021610) to Adrian Goldman, the Academy of Finland (No. 308105) to Keni Vidilaseris and (No. 310297) to Henri Xhaard and (279755) to Seppo Meri, and University of Helsinki funds to Gustav Boije af Gennäs. The authors thank Violeta Manole for her technical help, Nina Sipari for help with the MS-analyses, and Juho Kellosalo, Anssi Malinen and Peter Henderson for fruitful discussions during the project.

## Author Contributions

Conceptualization, K.V., A.G., H.X., J.Y.-K., G.B.G; Methodology, K.V., A.G., A.T., A.K., N.G.J., T.O.L.; Investigation, K.V., A. K., A.T, Ay.K.; Resources, A.G, H.X, S.M., J.Y-K.; Writing – Original draft, K.V., A.T., H.X., A.K; Writing-Review and Editing, K.V., A.K, A.T., Ay.K., N.G.J., T.O.L., P.S.K., G.B.G, S.M., J.Y-K., H.X, A.G.; Visualization, K.V., A.T., Ay.K.; Supervision and Project Administration, A.G., H.X, G.B.G, J.Y-K, S.M.; Funding Acquisition, A.G., H.X, K.V, J.Y-K., G.B.G, S.M.

## Accession codes

The atomic coordinates and structure factors of the TmPPase:IDP:ATC complex have been deposited in the Protein Data Bank, www.rcsb.org (PDB ID ^∗∗∗∗^).

